# Controlling for the effect of arterial-CO_2_ fluctuations in resting-state fMRI: Comparing end-tidal CO_2_ clamping and retroactive CO_2_ correction

**DOI:** 10.1101/2020.02.12.945881

**Authors:** Ali M. Golestani, J. Jean Chen

## Abstract

The BOLD signal, as the basis of functional MRI, arises from both neuronal and vascular factors, with their respective contributions to resting state-fMRI still unknown. Among the factors contributing to “physiological noise”, dynamic arterial CO_2_ fluctuations constitutes the strongest and the most widespread modulator of the grey-matter rs-fMRI signal. Some important questions are: (1) if we were able to clamp arterial CO_2_ such that fluctuations are removed, what would happen to rs-fMRI measures? (2) falling short of that, is it possible to retroactively correct for CO_2_ effects with equivalent outcome? In this study 13 healthy subjects underwent two rs-fMRI acquisition: During the “clamped” run, end-tidal CO_2_ (PETCO_2_) is clamped to the average PETCO_2_ level of each participant, while during the “free-breathing” run, the PETCO_2_ level is passively monitored but not controlled. PETCO_2_ correction was applied to the free-breathing data by convolving PETCO_2_ with its BOLD response function, and then regressing out the result. We computed the BOLD resting-state fluctuation amplitude (RSFA), as well as seed-independent mean functional connectivity (FC) as the weighted global brain connectivity (wGBC). Furthermore, connectivity between conditions were compared using coupled intrinsic-connectivity distribution (ICD) method. We ensured that PETCO_2_ clamping did not significantly alter heart-beat and respiratory variation. We found that neither PETCO_2_ clamping nor correction produced significant change in RSFA and wGBC. In terms of the ICD, PETCO_2_ clamping and correction both reduced FC strength in the majority of grey matter regions, although the effect of PETCO_2_ correction is considerably smaller than the effect of PETCO_2_ clamping. Furthermore, while PETCO_2_ clamping reduced inter-subject variability in FC, PETCO_2_ correction increased the variability. Overall PETCO_2_ correction is not the equivalent of PETCO_2_ clamping, although it shifts FC values towards the same direction as clamping does.

## Introduction

Resting-state functional magnetic resonance imaging (rs-fMRI) (Biswal et al., 1995) is a family of techniques whose role is rapidly expanding in neuroscience and clinical research (Biswal, 2012; Bookheimer et al., 2019; Liu et al., 2009). Despite the widespread adoption of rs-fMRI and associated functional connectivity (FC) measures in neuroimaging research, its meaningful interpretation is hampered by a lack of understanding of its biophysical mechanisms1. The blood-oxygenation level dependent (BOLD) signal, which is the basis of rs-fMRI, is an indirect measure of neuronal activity. The BOLD signal arises from both neuronal and vascular factors, with their respective contributions to rs-fMRI still unknown. Importantly, in aging and disease, interactions between these variables are often altered, making it extremely challenging to accurately interpret rs-fMRI results.

The impact of systemic physiological processes on rs-fMRI are an important open question. The most substantial contributors to the rs-fMRI signal are thought to be respiratory-volume variability, cardiac rate variability and carbon dioxide (CO_2_) fluctuations. While these processes are assumed to be unrelated to neuronal activity (Birn et al., 2006; Chang and Glover, 2009; Shmueli et al., 2007; Wise et al., 2004), they are found in the same frequency range as FC calculations (0.001 - 0.1 Hz), and are typically not explicitly removed before FC calculations (Smith et al., 2013). Specifically, the effect of CO_2_ was the first among the three to be investigated in rs-fMRI (Wise et al., 2004), and our previous work has demonstrated that dynamic CO_2_ fluctuations constitutes the strongest modulator of the rs-fMRI signal (Golestani et al., 2015), accounting for up to 15% of total rs-fMRI signal variance. Specifically, CO_2_ fluctuations are most strongly associated with rs-fMRI signals in the grey matter, notably in the temporal and occipital cortices as well as the thalamus, the superior parietal lobule, precuneus and cingulate gyrus (Golestani et al., 2015).

In terms of FC measurements using rs-fMRI, the study by Madjar et al. (Madjar et al., 2012) was the first to investigate the effect of eliminating PETCO_2_ fluctuations at the source through computerized PETCO_2_ targeting (i.e. PETCO_2_ clamping). It was found that clamping resulted in reduced resting FC in the default-mode network (DMN). However, a few questions remain unanswered: (1) it is unclear how PETCO_2_ clamping affects connectivity beyond the DMN; (2) it is unclear why no significant difference in percent BOLD fluctuations was found between free-breathing (FB) and clamped conditions; (3) it is unclear why an increased FC between DMN and occipital areas was observed under clamping. As this was the only relevant study to date, it is important to determine whether these results are reproducible on a broader scale. By studying the effect of CO_2_ clamping on rs-fMRI-based FC measurements, we expect to arrive at less ambiguous and more compelling conclusions regarding the effect of PETCO_2_ fluctuations on rs-fMRI.

Moreover, we pioneered the retroactive correction of PETCO_2_ fluctuation effects in rs-fMRI using voxel-wise estimation of the PETCO_2_ hemodynamic response function (HRF). In our previous work (Golestani et al., 2015), we parameterized a gamma-function form of a global-averaged PETCO_2_ HRF, which can facilitate the correction of PETCO_2_ effects when voxel-wise HRFs are not available. However, a natural question remains as to whether retroactive correction is equivalent to PETCO_2_ clamping.

In the current study, we compare rs-fMRI signals and FC measurements derived from (1) FB; (2) clamped and (3) retroactive PETCO_2_ corrected rs-fMRI data. In order to achieve clamping, we used the RespirAct system for prospective targeting of PETCO_2_. We hypothesize that (1) PETCO_2_ clamping should reduce the rs-fMRI signal fluctuation amplitude; (2) PETCO_2_ clamping should reduce the rs-fMRI FC values; (3) PETCO_2_ clamping should reduce the inter-subject variability in rs-fMRI FC measurements; (4) PETCO_2_ correction will produce subdued effects that are similar in nature to those of PETCO_2_ clampling.

## Materials and methods

### Participants and data collection

13 healthy subjects (6 males, age = 26.5 ± 6.5 years) were scanned using a Siemens TIM Trio 3T MRI scanner with a 32-channel head coil. Resting-state BOLD scans were collected using the simultaneous multi-slice GE-EPI BOLD technique (TR/TE = 380/30 ms, flip angle = 40°, 20 5-mm slices, 64×64 matrix, 4×4×5 mm voxels, multiband factor = 3, a total of 950 volumes) with leak-block slice GRAPPA recon with a 3×3 kernel (Cauley et al., 2014). For each subject, two resting-state fMRI dataset was collected, named the “clamped” and “free-breathing” runs. During the “clamped” run, we clamped PETCO_2_ to the average PETCO_2_ level of each participant using the RespirAct™ breathing circuit (Thornhill Research, Toronto, Canada). Subjects were instructed to breathe freely during the task while monitoring their air bags. For the “free-breathing” run, the PETCO_2_ level is passively monitored but not controlled. A T1-weighted anatomical image was collected for anatomical-registration purposes (MPRAGE, TR = 2400ms, TE = 2.43 ms, FOV = 256 mm, voxel size = 1×1×1 mm^3^).

### Physiological signals

During the fMRI scans, partial CO_2_ pressure (PCO_2_) fluctuations were recorded using a RespirAct™ breathing circuit (Thornhill Research, Toronto, Canada). In addition, cardiac pulsation was recorded using the Siemens scanner pulse oximeter (sampling rate = 50 Hz), whereas the respiratory signal was recorded using a pressure-sensitive belt connected to the BiopacTM (Biopac Systems Inc. California) at a sampling rate of 200 Hz.

The cardiac and respiratory signals were low-pass filtered to 2 Hz and 1 Hz, respectively using a Butterworth infinite-impulse response (IIR) filter with the corresponding bandwidths. Then, each signal is normalized by subtracting the mean and dividing by the standard deviation. Cardiac and respiratory frequencies are estimated as the peak frequencies of the spectra of the filtered (infinite-impulse response) and normalized cardiac and respiratory signals, respectively.

In addition to original physiological signals, low-frequency variations of the physiological signals were calculated. Cardiac-rate variation (CRV) is calculated as the average heart-beat rate over a 4 second window (change et al 2009). Respiratory-volume variability (RVT) is defined as the ratio of breathing dept (difference between maximum and minimum) over breathing period (time elapsed between consecutive maxima) (Birn et al 2006). PETCO_2_ is calculated by finding the peak PCO_2_ level in each breathing cycle (Golestani et al 2015), and outliers beyond three standard deviations of each time course were replaced with interpolated values based on neighbouring points. Subsequently, the mean and standard deviation of these physiological signals were calculated.

### fMRI analysis

The rs-fMRI processing pipeline includes: motion correction, spatial smoothing (Gaussian kernel with 5mm FWHM), high-pass filtering (> 0.01 Hz), and registration of data into a 4×4×4 mm^3^ MNI atlas (45 × 54 × 45 voxels). All steps were performed using FMRIB software library (FSL, publicly available at www.fmrib.ox.ac.uk/fsl). We assessed bulk-head motion in terms of framewise displacement (FD).

To provide a seed-independent assessment of mean FC strength, we used the weighted global brain connectivity (wGBC) measure (Cole et al., 2010; Scheinost et al., 2014, 2012). We first define ***f*(*x*, *r*)** as the distribution of correlations (r) for voxel ***x***, where ***r*** is the correlation coefficient relating the time series in voxel x to that of another voxel in the gray matter. We can thus construct a correlation map for voxel x for each condition. From this, we can compute the wGBC (Cole et al., 2010), which is the average of all Fisher-transformed r values within the distribution. A higher wGBC indicates higher global connectedness. We also computed the resting-state fluctuation amplitude (RSFA) for each subject in each condition.

### Retroactive PETCO_2_ correction

In our previous research we modeled the effect of the PETCO_2_ signal on the fMRI signal and proposed a voxel-wise method for correcting the effect of the PETCO_2_ fluctuations by regressing out its effect from the BOLD signal. To investigate if such a correction can eliminate the effect of PETCO_2_ fluctuation,the effect of PETCO_2_ fluctuation is removed from free-breathing data by voxel-wise estimating response function of the BOLD signal to PETCO_2_ signal. We followed the procedure outlined in our previous work. First, the PETCO_2_ time course was orthogonalized against the RVT and CRV time courses. Then the result is deconvolved from the voxel-wise BOLD fMRI signal to obtain the PETCO_2_ response function. The PETCO_2_ is then convolved with the response function and regressed out of the BOLD signal at each voxel, producing the corrected free-breathing data. Refer to (Golestani et al., 2015) for more details. We then compared the connectivity strength and variability between the clamped and corrected free-breathing data.

### Statistical testing

#### Effect of clamping on physiological signals

Although the clamped PETCO_2_ condition was designed to target each subject’s true resting-state PETCO_2_, the PETCO_2_ level climbed slightly during each clamped session (see Table 1). Therefore, we regressed out the effect of individual mean-PETCO_2_ levels before assessing the difference between clamped and free-breathing results.

**Table 1.** Comparison of physiological signal features and estimated head motion between free-breathing and clamped conditions using paired t-test. The values are given in the format of mean ± stdev computed across all subjects.

|  |  | Free breathing | Clamped | Significance of difference (p value) |
| --- | --- | --- | --- | --- |
| Cardiac frequency (Hz) | | 1.11 ± 0.21 | 1.11 ± 0.20 | 0.92 |
| Respiration frequency (hz) | | 0.23 ± 0.05 | 0.23 ± 0.069 | 0.92 |
| CRV | Mean | 0.93 ± 0.16 | 0.92 ± 0.159 | 0.38 |
| | Stdev | 0.0049 ± 0.016 | 0.029 ± 0.012 | **0.0002** |
| RVT | Mean | 0.0046 ± 0.0037 | 0.0050 ± 0.0026 | 0.63 |
| | Stdev | 0.0013 ± 0.0012 | 0.0014 ± 0.0016 | 0.68 |
| PETCO<sub>2</sub> | Mean | 37.68 ± 2.60 | 40.69 ± 2.20 | **1.02e-5** |
| | Stdev | 1.0020 ± 0.25 | 0.69 ± 0.33 | **0.001** |
| Framewise displacement | Mean | 0.067 ± 0.019 | 0.067 ± 0.020 | 0.94 |
| | Stdev | 0.038 ± 0.013 | 0.037 ± 0.013 | 0.87 |

Cardiac and respiratory signals across multiple subjects were compared between free-breathing and clamped conditions using paired t-test. To do this, the spectra of clamped cardiac and respiratory signals were subtracted from those of the free-breathing condition for each individual. The difference spectra were then tested for statistical significance at frequency intervals of 0.0056 Hz for all frequencies in range (0 to 2 Hz for cardiac and 0 to 1 Hz for respiratory signals). The frequencies at which the difference spectra are significantly different from zero are identified (p < 0.05, false-discovery rate corrected).

The spectra of PETCO_2_, RVT and CRV were also compared between free-breathing and clamped conditions using paired t-tests, sampling over a frequency range of 0 to 1.3 Hz at steps of 0.0033 Hz. As before, we used false-discovery rate (FDR) for multiple-comparisons correction.

#### Effect on rs-fMRI signal and functional connectivity

First, we compared wGBC and RSFA across pairs of conditions at a group level using paired t-tests, implemented using Matlab (Mathworks, Natick, USA).

To assess the difference in connectivity strength between pairs of conditions, we used the coupled intrinsic connectivity distribution (ICD) method (Scheinost et al., 2014), which is designed to compare functional connectivity maps between two conditions in the same cohort. To briefly summarize the method, the coupled ICD method is based on the ICD metric (Scheinost et al., 2012), which is derived by parameterizing the survival function of f(x, r) at a given threshold τ.

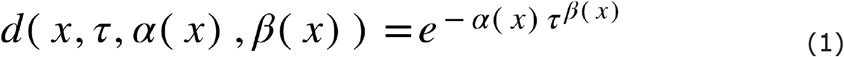

In coupled ICD, instead of defining a survival function d(.) at each voxel, we subtract the correlation distribution ***f*(*x*, *r*)** in one condition from that of another condition, and the resulting distribution of ***r*** differences is modeled as a Weibull distribution with a variance parameter *α* and slope parameter *β*:

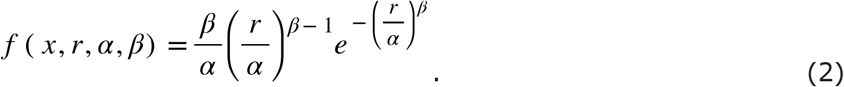

This expression models the distribution and variance of r for each voxel in terms of *α* (the variance parameter) and *β* (the shape parameter, also known as the Weibull slope). A larger α value indicates a greater variance in the difference distribution, conveying that a relatively high number of correlations have changed between conditions. On the other hand, a higher β value indicates the slope of the tail (left or right) of the distribution. These parameters are then submitted to statistical comparisons through paired t-tests. Specifically, when the ***f*(*x*, *r*)** for each condition is subtracted from that from another condition, positive and negative differences can result, associated with the right and left side of the Weibull distribution, respectively.

The coupled-ICD approach provides (1) a more comprehensive description of differences between conditions than conventional methods, representing the r distribution at each seed voxel as a distribution rather than solely as a mean value (wGBC); (2) a straight-forward way to perform statistical testing, by simply comparing the distribution of *α* and *β* values (one distribution for each subject). When the f(x, r) map of condition A is subtracted from that of condition B, the Weibull distribution is fitted to positive (right tail) and negative values (left tail) of the connectivity-difference distribution separately. To compare the connectivity strength between the two conditions, *α* and *β* values were compared between left and right tails across subject, using paired t-test. The analysis steps are summarized in **Figure 1**.

**Figure 1.**
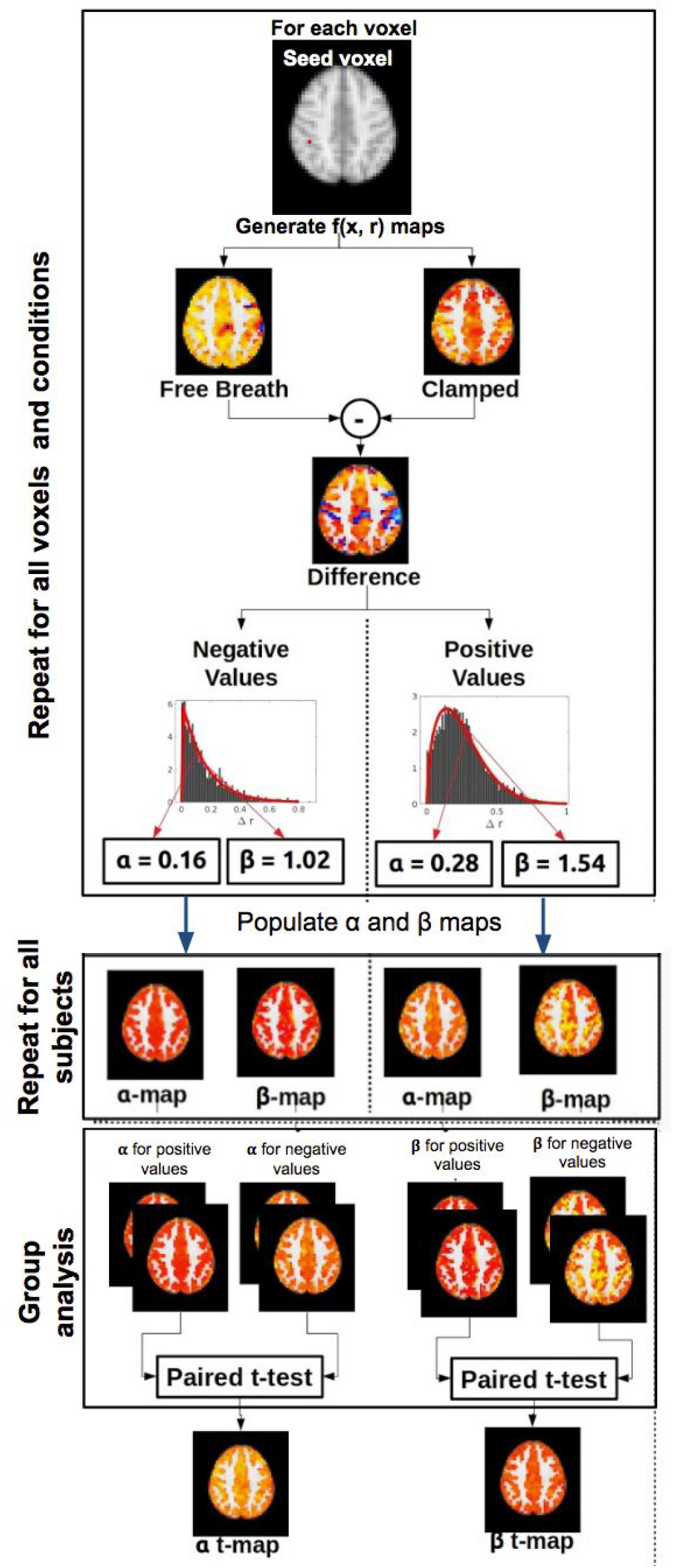
Summary of analysis steps for comparing connectivity strength across conditions using the coupled-ICD approach. In this case, the example of “free-breathing vs. clamped” is shown. At each voxel, the distribution of individual between-condition differences in FC is generated by subtracting ***f*(*x*, *r*)** of one condition from that of another condition (e.g. clamped from free-breathing in this case). We use these difference maps to produce distribution of positive and negative differences, each characterized by its **α** and **β** parameters. The associated parametric maps of the left (negative difference) and right (positive difference) distributions are compared to generate the t maps for these parameters.

In the coupled-ICD framework, if the connectivity in the two conditions are similar, the distribution should be symmetrically distributed around zero and therefore *α* and *β* should be similar for left and right tails. Conversely, if the *α* value in the positive half of the difference distribution is significantly greater than that of the negative tail, we conclude that FC distribution include higher values in condition B than in condition A (Scheinost et al., 2014). Likewise, if the *β* value in the positive half of the difference distribution is significantly greater than that of the negative tail, we may conclude that mean of the FC distribution is higher in condition B than in condition A.

We further compared the inter-subject variability in the connectivity values. Again, for each seed voxel from each data set (subject), an r map is generated. Assessing the variability of these r maps across subjects provides the inter-subject variability map for each condition. In each variability map, outliers beyond the 95% confidence interval of the distribution are replaced by the median. This variability map is then compared between conditions, and a significance value determined from a paired z-test. The z-test is chosen due to the large sample size (12,000 voxels in each FC map). We then repeated this process for each seed voxel to assess the significance for the difference in inter-subject variability between conditions. The steps are summarized in **Figure 2**. For voxels reaching statistical significance (p < 0.05, corrected for multiple comparisons) in terms of the inter-condition difference, we computed the effect size (i.e. difference in inter-subject variability across conditions) using Cohen’s D, computed as |μ1 − μ2|/σ, where μ_1_ and μ_2_ are the means from the two conditions, and σ is the square root of the average variance in each condition (assuming equal sample sizes in both conditions).

**Figure 2.**
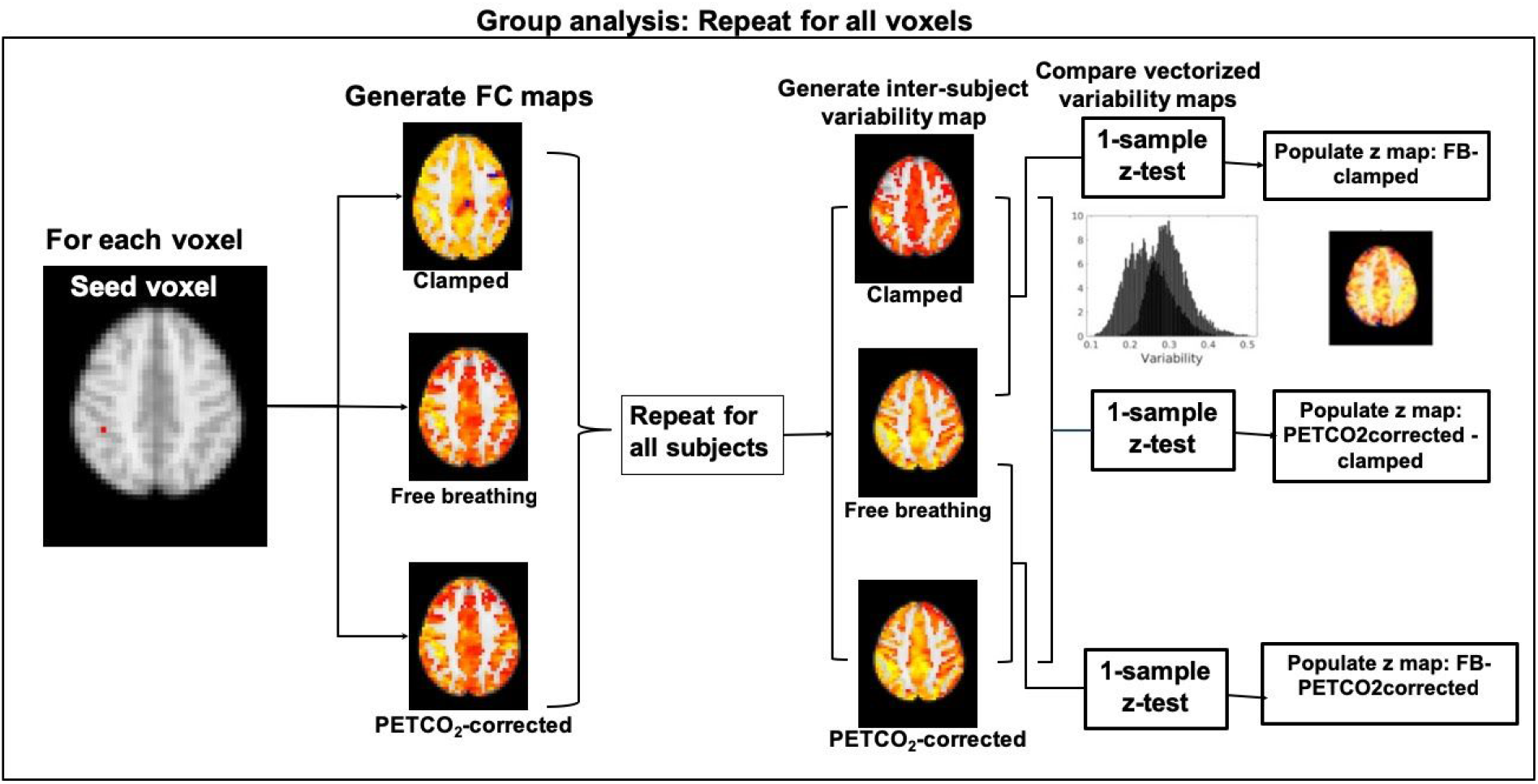
Summary of analysis steps for comparing FC inter-subject variability across conditions. Each grey matter voxel is considered as a seed and its associated connectivity maps are generated for all subjects. We further assess the inter-subject variability in these r maps, generating an inter-subject variability map for each condition (namely, free-breathing, clamped or PETCO_2_-corrected). This variability map is then compared between conditions, and a significance value determined from a paired z-test.

Finally, all statistical results, which were already in MNI152 space, were mapped to the FreeSurfer average brain surface (fsaverage) using FreeSurfer 6.0 (publicly available: https://surfer.nmr.mgh.harvard.edu).

## Results

### Effect of clamping on cardiac and respiratory signals

We found cardiac and respiratory frequencies not to be significantly different between clamped and free-breathing conditions (Figure 3, top panel), amidst much inter-subject variability in the differences. Moreover, the spectral comparisons show no significant difference in the cardiac and respiration spectra between the two conditions (Figure 3, bottom panel). Furthermore, head motion, as quantified by framewise displacement (FD), are also not significantly different between the two conditions (Table 1).

**Figure 3.**
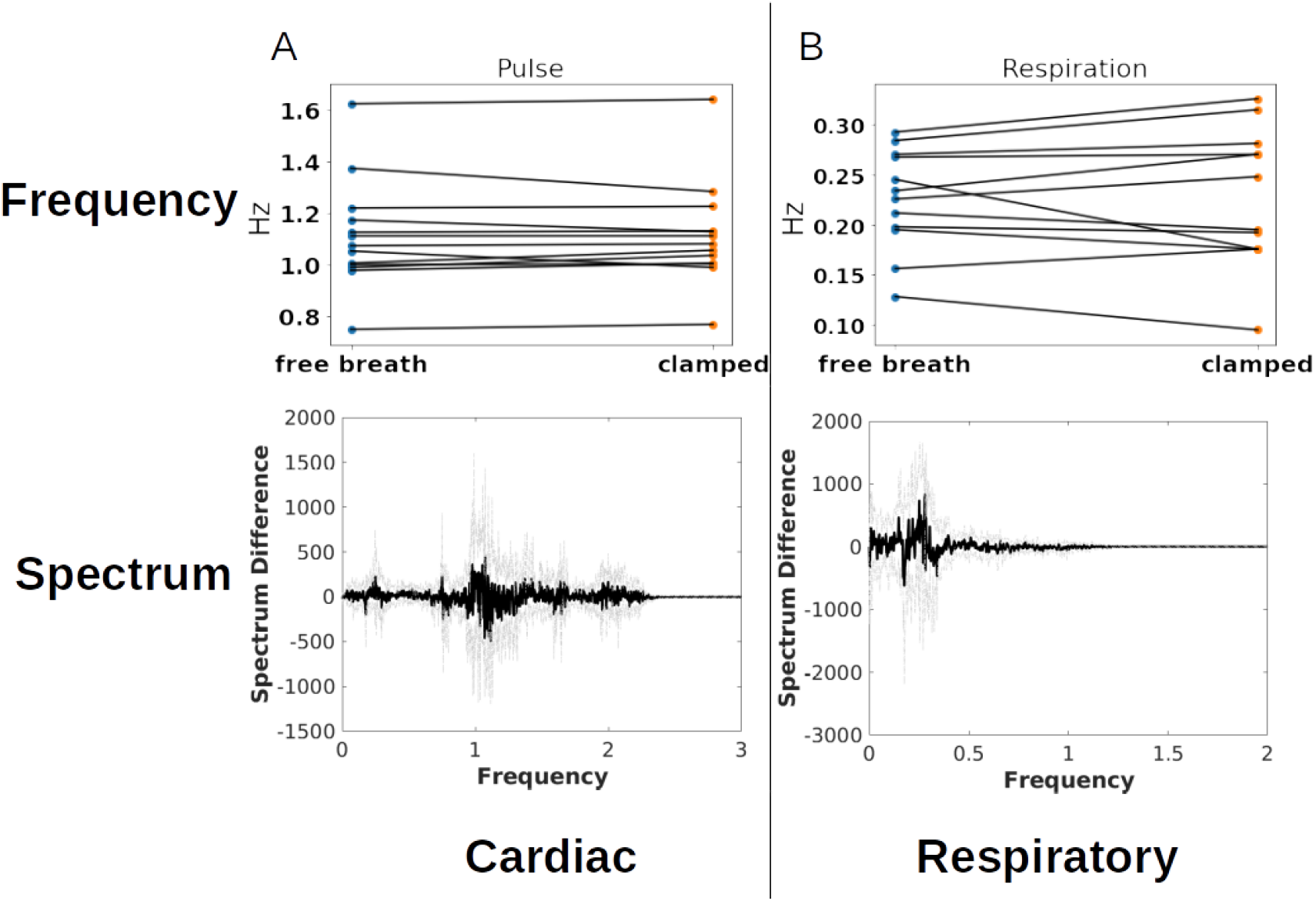
The effect of PETCO_2_ clamping on the cardiac (A) and respiratory signals (B). Slope-charts of heart rate (A) and respiration rate (B) are shown in the top row to illustrate inter-subject variability in the between-condition differences. Group-wise paired t-test showed no difference between conditions in terms of heart rate and respiration rate. The difference between clamped and free-breathing power spectra of cardiac and respiratory signals are shown in the bottom panel. The dark line represents the group mean, and the dotted lines represent the group standard deviation in the difference spectra. The frequency spectra not significantly different between the free-breathing and clamped conditions.

### Effect of clamping on PETCO_2_, CRV and RVT

While the mean CRV values from the free-breathing and clamped conditions are similar (p = 0.38), the temporal standard deviation of the mean CRV is significantly reduced in the clamping condition (p = 0.0002). Comparing the CRV spectra (at intervals of 0.0033 Hz) between conditions also showed significant difference at ∼ 0.13Hz (Figure 4G, bottom plot). The average and standard deviation of RVT are not significantly different between free-breathing and clamped conditions (p = 0.63). However, the standard deviation of the PETCO_2_ time series is significantly lower in clamped condition (p = 0.001). This is expected, as the goal of clamping is reducing PETCO_2_ fluctuations. As mentioned earlier, although we tried to match the level of PETCO_2_ in the clamped condition to the average level of PETCO_2_ in free-breathing for each individual, the average level of PETCO_2_ is slightly but significantly higher in the clamped condition compared to the free-breathing (p = 1e-5, see Table 1). Therefore the level of PETCO_2_ is regressed out in all future analysis. No significant difference in the spectra of PETCO_2_ is observed between the two conditions at any specific frequency.

**Figure 4.**
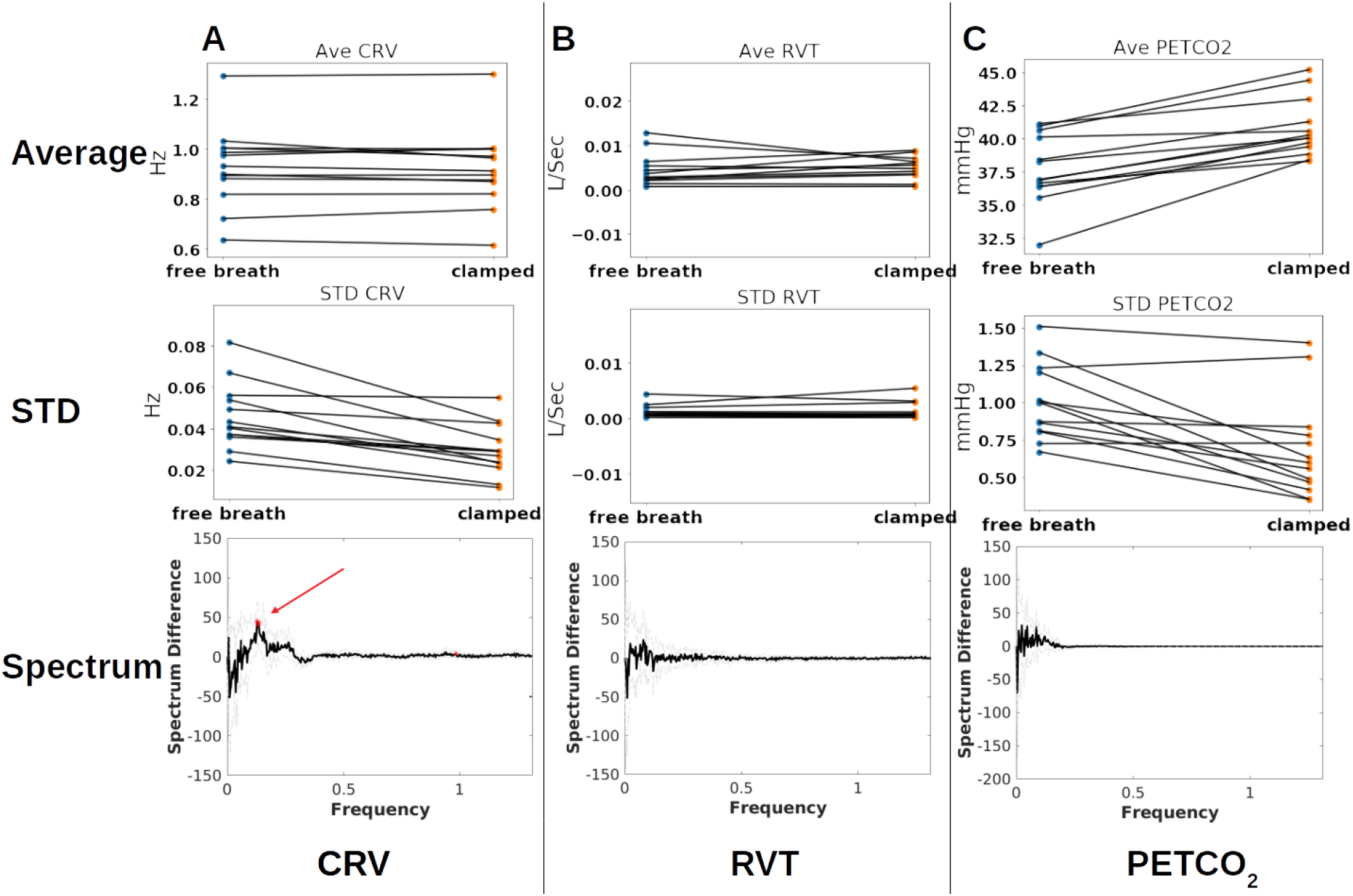
The effect of PETCO_2_ clamping on CRV (A) RVT (B) and PETCO_2_ fluctuations (C). Top panel shows comparison of the average values, middle panel shows comparison of the standard deviations, and bottom panel shows the spectrum comparison. Clamping PETCO_2_ resulted in reduced standard deviation in CRV and spectrum alteration around 0.13 Hz. RVT is not different between the two conditions. PETCO_2_ clamping did not change the PETCO_2_ spectrum, yet it significantly reduced the standard deviation of PETCO_2_ fluctuations. Moreover, average PETCO_2_ level is significantly higher in clamped condition compared to free-breathing.

### Effect of PETCO_2_ clamping and correction on RSFA and FC

As seen in Figure 3(a), when summarized across all voxels and subjects, the group difference in RSFA between free-breathing (FB) and clamped conditions exceeds that of the difference between CO_2_-corrected and clamped. The same trend is observed in the case of global mean FC (wGBC), but the difference between free-breathing and CO_2_-corrected conditions is greater than in the RSFA case. As shown in Figure 3 (c) and (d), the difference in RSFA and wGBC values between CO_2_-corrected and clamped is smaller than the difference between FB and clamped at the individual-subject level for some subjects. However, across the group, the RSFA and wGBC differences between conditions was not significant.

### Effect of PETCO_2_ clamping and correction on intrinsic connectivity distribution

In terms of the ICD, the outcome of free-breathing minus clamped conditions is associated with significantly greater α values for the positive tail than for the negative tail of the difference distribution. Spatially, this effect is associated with most cortical regions (Figure 6a). This finding is further illustrated by comparing the green histograms in Fig. 7 (a) and (c). A similar trend is observed in β, but affecting a smaller collection of cortical regions (Fig. 6b), and illustrated by the green histograms in Fig. 7 (b) and (d).

**Figure 6.**
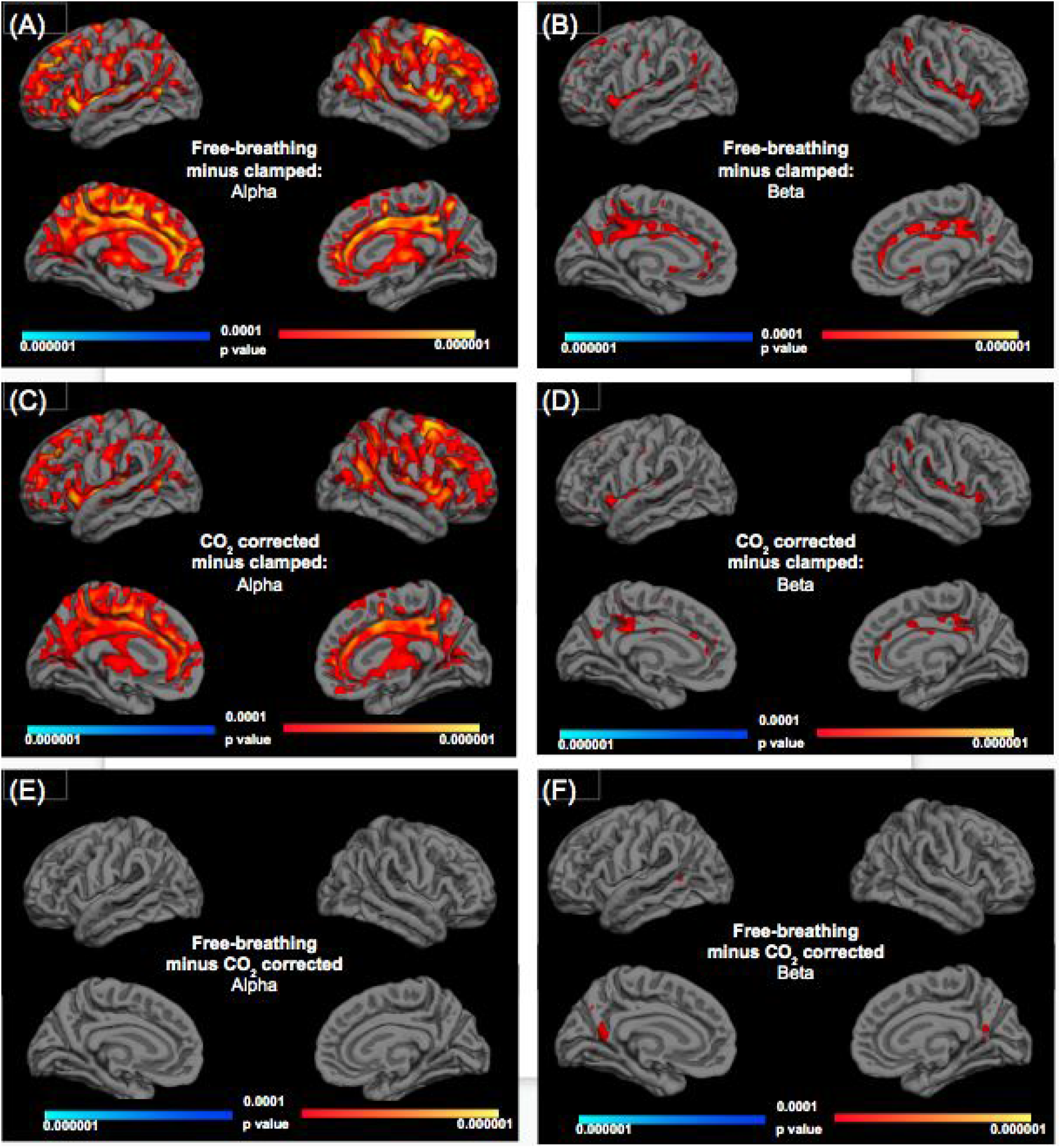
The difference in FC maps across the three conditions. We show differences in wGBC distributions, summarized by **α** and **β**, which represent the variance and the slope (of the tail) of the difference distribution (computed between conditions). In each instance, statistically significant effects are shown for the lateral (top panel) and medial (bottom panel) aspects of the cortex for both left and right hemispheres. Orange and blue indicate positive and negative **α** and **β** effects, respectively. The strongest effects are found in **α** and when subtracting “clamped” from “free-breathing” conditions, peaking in the superior precentral gyrus, the precuneus, the temporoparietal junction, the insula, the superior frontal cortex and the entire cingulate cortex (A). We also noted particularly strong effects in the thalamus. This is similar to the pattern exhibited by the “CO_2_-corrected minus clamped” case (C). The effects in **β** are much weaker and more sparse (B, D). Finally, minimal differences were found between CO_2_-corrected and free-breathing conditions for either **α** or **β**.

**Figure 7.**
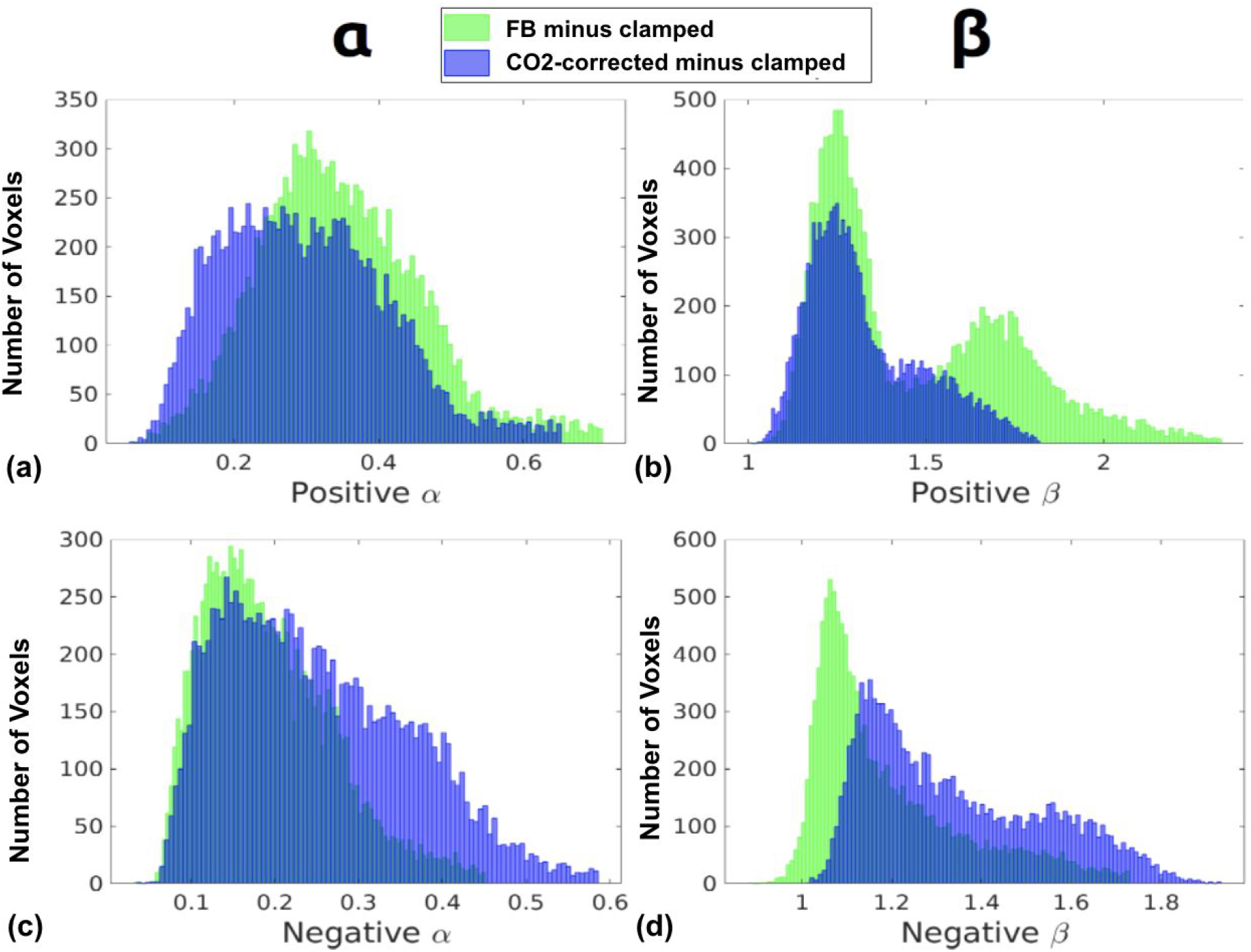
**Histograms of FC differences between conditions**, summarizing all voxels from a representative subject. Positive and negative **α** refer to **α** values extracted from the distributions of positive and negative tails of the Weibull distribution, respectively. The same applies to positive and negative **β**. For instance, in the case of “free-breathing (FB) minus clamped”, higher **α** values are observed than in the case of “CO_2_-corrected minus clamped” (A), indicating that FB without CO_2_ correction is generally associated with greater FC variance than in the CO_2_-corrected case. The latter, in turn is associated with greater FC variance than the clamped case. The strongest effects for **α** are found in the superior precentral gyrus, the precuneus, the temporoparietal junction, the insula, the superior frontal cortex, the entire cingulate cortex and part of the thalamus (A). This is similar to the pattern exhibited by the “CO_2_-corrected minus clamped” case (C). Likewise, a higher **β** in the “free-breathing (FB) minus clamped” case compared to the “CO_2_-corrected minus clamped) case indicates that the former difference distribution is associated with a shift in the mean FC towards a higher value.. The effects in **β** are much weaker and more sparse (B, D). Finally, minimal differences were found between CO_2_-corrected and free-breathing conditions for either **α** or **β**.

Interestingly, in the case of α, the application of PETCO_2_ correction did not result in a substantial difference compared to free-breathing. That is, when taking the difference in f(x, r) between CO_2_-corrected and clamped conditions (Fig. 6(c)), the voxels exhibiting significantly different α values between the positive and negative tails of the difference distribution visually overlap almost completely with the voxels in Fig. 6(a), although the statistical significance is lower over all. The same trend is observed for β (Fig. 6(d)).

To clarify the changes underlying Fig. 6, we plotted the α and β histograms associated with the positive and negative difference distributions for a representative subject in Fig. 7. As shown in Fig. 7b, correction for the PETCO_2_ fluctuation reduced the number of voxels with significantly higher α. However, PETCO_2_ correction enlarged the distance between the positive β and negative β distributions, as seen by comparing the blue and green histograms in Fig. 7b and 7d.

### Effect of PETCO_2_ clamping and correction on FC inter-subject variability

In Figure 8 are shown the difference in the inter-subject variability in functional connectivity (FC) values between clamped and free-breathing conditions. PETCO_2_ clamping results in significantly lower between-subject variability in FC compared to free-breathing data in almost the majority of grey matter regions (Fig. 8a). The same is true when comparing clamped to CO_2_-corrected conditions. However, when compared to the free-breathing condition, CO_2_ correction resulted in higher inter-subject FC variability (Fig. 8b). Across all three figures, the strongest effects (positive or negative) are found in the insula, the precuneus, the cingulate cortex, the superior parietal lobule and the temporoparietal junction. While we do not delve deeply into the deep-grey structures or the white matter, we noted particularly strong effects in the thalamus once again.

**Figure 8.**
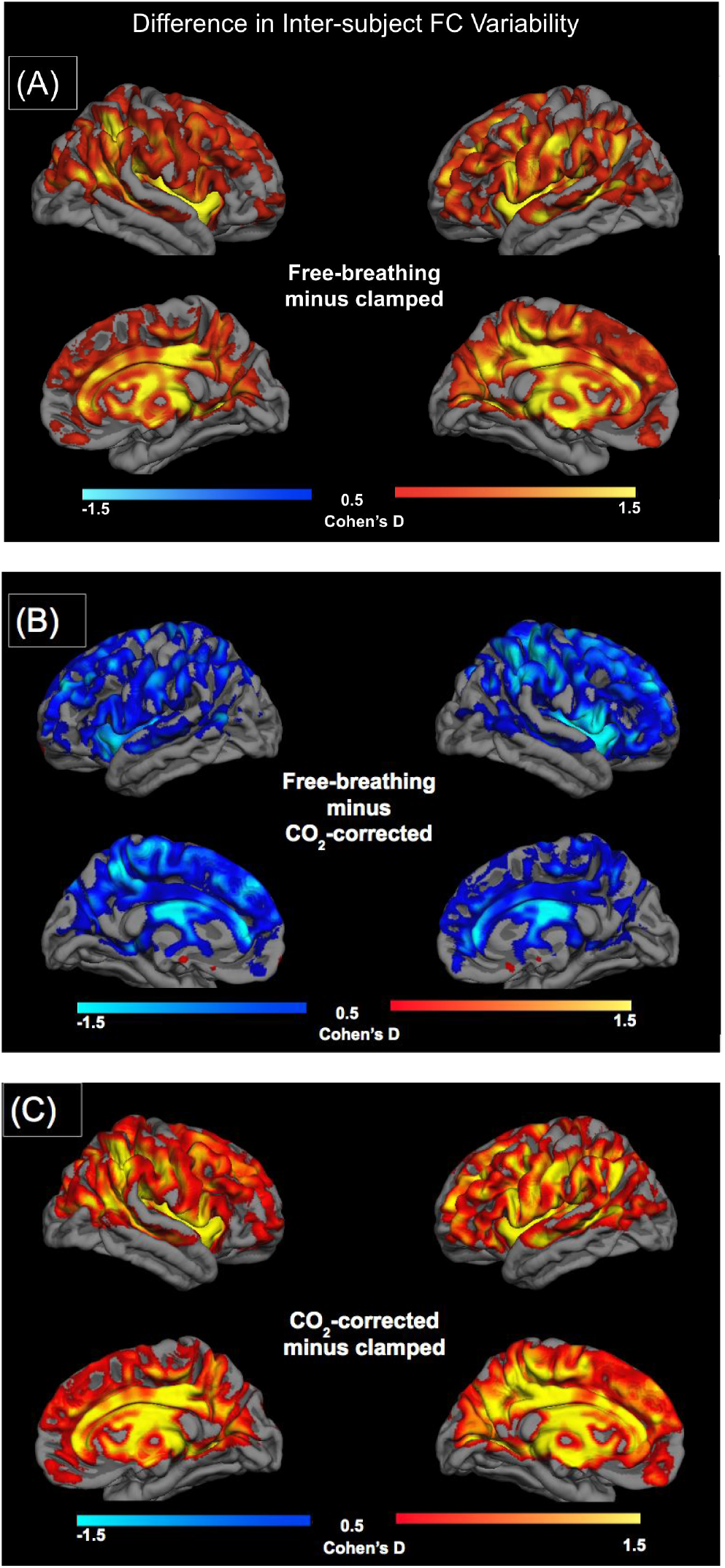
Cross-condition differences in inter-subject variability of functional connectivity. Free breathing is associated with significantly higher inter-subject FC variability compared to clamped (A), whereas CO_2_-correction appears to result in an inter-subject FC variability that is higher than free breathing (B) as well as than clamped (c). In each instance, statistically significant effects are shown for the lateral (top panel) and medial (bottom panel) aspects of the cortex for both left and right hemispheres. Orange and blue indicate positive and negative differences, respectively. Across all three figures, the strongest effects are found in the insula, the precuneus, the cingulate cortex, the superior parietal lobule and the temporoparietal junction. The thalamus also showed significant effects over all.

## Discussion

In this work, we quantify the difference between rs-fMRI measures captured during PETCO_2_ clamping, free breathing and free breathing after PETCO_2_ correction. The main findings are: (1) PETCO_2_ clamping did not significantly alter RVT and CRV; (2) PETCO_2_ clamping resulted in a significant reduction in the functional connectivity (FC) as computed using voxel seeds; (3) PETCO_2_ clamping reduced Inter-subject variability in functional connectivity; (4) PETCO_2_ correction increased inter-subject variability in FC. We discuss each finding as follows.

Sustained increases in arterial CO_2_ is known to suppress neuronal activity (Driver et al., 2016; Hall et al., 2011; Uh et al., 2009), as measured using electroencephalography (Xu et al., 2011) and magnetoencephalography (Driver et al., 2016; Hall et al., 2011). This observation spans the delta, alpha, beta and gamma bands. Notably, in the resting state, CO_2_ fluctuations accounts for 2% of MEG activity variance (r = 0.15) (Driver et al., 2016). The main mechanism is proposed to be pH, which is inversely correlated with extracellular adenosine concentration. Elevated extracellular CO_2_ reduces neuronal excitability, but is not likely to overcome the intracellular pH buffering to affect intracellular pH. As CO_2_ fluctuations are part of respiration, its effect on rs-fMRI is presumably not functional specific.

It is well established that fluctuations of arterial CO_2_, measured indirectly through end-tidal CO_2_ (PETCO_2_), constitutes a sizeable source of rs-fMRI signal fluctuation (Wise et al., 2004). The resting-state BOLD signal is most sensitive to PETCO_2_ fluctuations in regions with high vascular density or high metabolic activity levels (Golestani et al., 2015; Wise et al., 2004). Wise et al. found the effect to peak in the superior parietal cortex, the paracentral cortex, the temporoparietal junction, the insula and the thalamus. Later, Golestani et al. independently found the peak PETCO_2_ effects on the rs-fMRI signal in the temporal and occipital cortices as well as the superior parietal lobule, precuneus, the cingulate gyrus and the thalamus (Golestani et al., 2015). Therefore, the literature is consistent on the regions of impact of PETCO_2_ fluctuations.

Currently the only ways to achieve PETCO_2_ clamping are to use end-tidal forcing (Wise et al., 2007) or prospective PETCO_2_ targeting (Prisman et al., 2008). Madjar et al. (Madjar et al., 2012) pioneered the use of PETCO_2_ clamping in fMRI using the latter approach, which is the only way to fix each subject’s PETCO_2_ and PETO_2_ to his/her natural baseline mean levels. They found clamping to result in reduced FC in the default-mode network and increased FC in the visual network. Our work adopts the same clamping approach as Madjar et al.. Our use of seed-independent connectivity and the coupled-ICD approach allowed us to observe such an effect in more regions than previously reported (Madjar et al., 2012). We found that PETCO_2_ clamping contributed to significant and widespread coupled-ICD differences, linking higher CO_2_ fluctuations to higher rs-fMRI functional connectivity (FC) measures across the cortex. The regions most strongly affected are the superior precentral gyrus, the precuneus, the temporoparietal junction, the insula, the superior frontal cortex, the entire cingulate cortex and part of the thalamus (Fig. 6A, C). These resemble those reported (Golestani et al., 2015), suggesting that PETCO_2_-induced variability in the rs-fMRI signal translate directly into higher FC. However, of the impacted regions in both cases, the cingulate, which exhibited FC reduction with clamping, did not exhibit PETCO_2_ effects on rs-fMRI amplitude. Conversely, the occipital cortex, which exhibited PETCO_2_ effects in its signal amplitude, was not found to exhibit PETCO_2_ effects on FC. The significance of this mismatch warrants further investigations.

As expected, PETCO_2_ clamping resulted in a reduction in inter-subject FC variability, as shown in Fig. 7. This is consistent with past work by Birn et al., who showed that physiological corrections generally reduced inter-subject variability (Birn et al., 2014). As inter-subject consistency is widely used as a quality metric for rs-fMRI measurements (Moussa et al., 2012; Ren et al., 2017), clamping presumably enhances the contribution of neuronally-specific inter-subject variability.

Given that clamping the resting-state arterial CO_2_ requires dedicated equipment with limited accessibility, in recent years the focus has been on correcting for this effect post hoc. Notably, we were the first to parameterize a CO_2_ response function (HRF_CO2_) as well as to show its spatial variability (Golestani et al., 2015). Recently, Prokopiou et al. presented an alternate method for HRF_CO2_ estimation, focusing on the difference between resting and task-driven HRF_CO2_ (Prokopiou et al., 2018). To retroactively correct for the effect of CO_2_ fluctuations on the fMRI signal, HRF_CO2_ is typically convolved with PETCO_2_ recordings, and the result regressed out of the fMRI signal. Finally, as seen in our results (Fig. 5), PETCO_2_ correction also reduced the RSFA and wGBC quite noticeably in certain subjects, although not significantly across the entire group of subjects.

**Figure 5.**
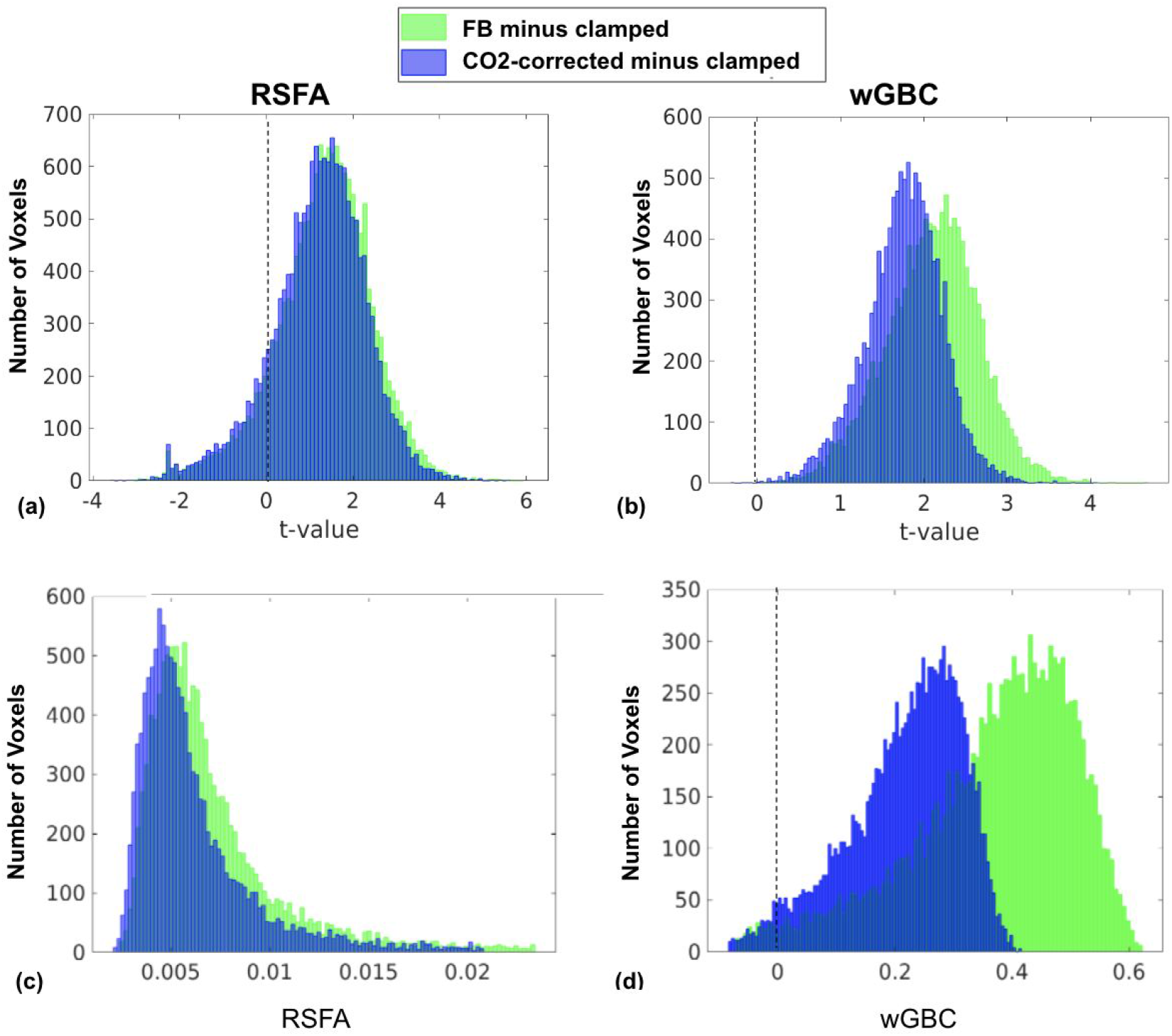
Histogram of inter-condition differences for RSFA and wGBC. (a) and (b) represent t-value distributions generated from comparing RSFA and wGBC between conditions. All voxels from all subjects are used in generating the histograms. The t-value distributions are more positive than negative, demonstrating that free breathing and CO_2_-correction are both associated with higher RSFA (a) and wGBC (b) values than clamped. However, the difference was not statistically significant once corrected for multiple comparisons. (c) and (d) represent the underlying data from a representative subject. Free-breathing (FB) is associated with higher RSFA values than clamped (green).

One of the motivations of our study is to compare the effects of post hoc PETCO_2_ correction against that of PETCO_2_ clamping in rs-fMRI. Contrary to our expectations, PETCO_2_ correction produced no significant change in group-level RSFA and wGBC, although it reduced RSFA and wGBC in several subjects. Furthermore, PETCO_2_ correction had only minimal effects on the coupled-ICD alpha measure (Fig. 6). Nonetheless, as reported in our previous work, PETCO_2_ correction improved the within-subject reproducibility of FC results (Golestani et al., 2017). It is unclear to us if improved within-subject reproducibility necessarily leads to increased inter-subject variability in FC. As Birn et al. reported, physiological corrections generally reduced within-subject variability, but also significantly reduced inter-subject variability (Birn et al., 2014). While Birn et al. did not examine the effect of PETCO_2_ per se, the finding of increased inter-subject variability with PETCO_2_ correction has interesting implications. As Birn et al. pointed out, physiological effects are reliable, so the removal thereof may in fact reduce the reliability of FC measurements. By the same token, the reduced reliability could lead to reduced inter-subject reproducibility. Nonetheless, we observed reduced inter-subject variability when PETCO_2_ was clamped, refuting this argument.

While these findings regarding PETCO_2_ correction contradict our hypotheses, they may be timely for the resulting questions regarding our assumptions regarding the brain’s response to CO_2_. Spontaneous fluctuations in PETCO_2_ are known to be involved in chemo-reflex-feedback regulation of subsequent breaths’ depth and rate (Modarreszadeh and Bruce, 1994; Van den Aardweg and Karemaker, 2002). This feedback cycle, lasting 25 s or longer, can induce changes in subsequent levels of arterial CO_2_. Regressing out PETCO_2_ fluctuations does not take into account this feedback cycle and therefore does not remove all BOLD signal fluctuations introduced by spontaneous fluctuations in PETCO_2_.

## Limitations

As mentioned earlier, while we managed to eliminate a large portion of the natural PETCO_2_ fluctuation by clamping, the mean clamped PETCO_2_ levels were slightly higher than the mean PETCO_2_ in the free-breathing state (Table 1). Nonetheless, the difference is small, and not likely to account for all of our findings. Moreover, the difference was factored into our statistical analyses (we regressed out mean-PETCO_2_ as a covariate in our general linear models).

Moreover, the metabolic impact of PETCO_2_ clamping was not explored in this study. Since the clamping target is determined by each subject’s natural baseline PETCO_2_, the only metabolic demand would stem from the respiratory task associated with prospective targeting. Nonetheless, as our subjects were instructed to breathe freely during the clamped session, the associated neuronal influence is expected to be minimal.

Finally, although we did not observe statistically significant changes in RSFA with PETCO_2_ correction or clamping, we believe the effect should be more significant in a larger sample. We aim to reproduce or confirm our findings on a larger scale in future work.

## Acknowledgments

This work has been supported by the Canadian Institutes of Health Research.

